# Long-term survival of salmon-attached SARS-CoV-2 at 4°C as a potential source of transmission in seafood markets

**DOI:** 10.1101/2020.09.06.284695

**Authors:** Manman Dai, Huanan Li, Nan Yan, Jinyu Huang, Li Zhao, Siqi Xu, Shibo Jiang, Chungen Pan, Ming Liao

## Abstract

Several outbreaks of COVID-19 were associated with seafood markets, raising concerns that fish-attached SARS-CoV-2 may exhibit prolonged survival in low-temperature environments. Here we showed that salmon-attached SARS-CoV-2 at 4°C could remain infectious for more than one week, suggesting that fish-attached SARS-CoV-2 may be a source of transmission.

## Text

The first outbreak of COVID-19 in late 2019 and early 2020 was associated with the Huanan Seafood Market in Wuhan, China, while the second outbreak of COVID-19 in June of 2020 was associated with the Xinfadi Seafood Market in Beijing, China (*1*–*2*). Several groups in different countries have identified SARS-CoV-2 in meat or meatpacking workers (*3*–*7*), raising concerns that fish- or meat-attached SARS-CoV-2 could be a potential source of COVID-19 transmission. Therefore, it is essential to determine the survival time of SARS-CoV-2 in the low-temperature environment of seafood markets.

In this study, we detected the titer (50% tissue culture infectious dose/mL, TCID_50_/mL) of viable SARS-CoV-2 attached on salmon or untreated SARS-CoV-2 in culture medium stored at 4°C, the temperature in refrigerators or cold rooms for the temporary storage of fish, or 25°C, the regular room temperature, respectively, using end-point titration assay on Vero E6 cells as described previously (*8*). As shown in Figure A and B, salmon-attached SARS-CoV-2 remained viable at 4°C and 25°C for 8 and 2 days, respectively, while the untreated SARS-CoV-2 in culture medium remained infectious at 4°C and 25°C for more than 8 days. SARS-CoV-2 attached on salmon or suspended in culture medium stored at 4°C remained viable for at least 8 days, while these stored at 25°C resulted in attenuating infectivity very quickly. The result from the experiment on samples stored at 25°C is consistent with that reported by van Doremalen et al. They showed that SARS-CoV-2 remained viable in aerosols, or on the surface of copper, cardboard, stainless steel, and plastic, at 21~23°C and 40% relative humidity for 3 ~ 24 hours (*8*), confirming that the loss of SARS-CoV-2 viability is associated with increased temperature.

**Fiugre A and B.**
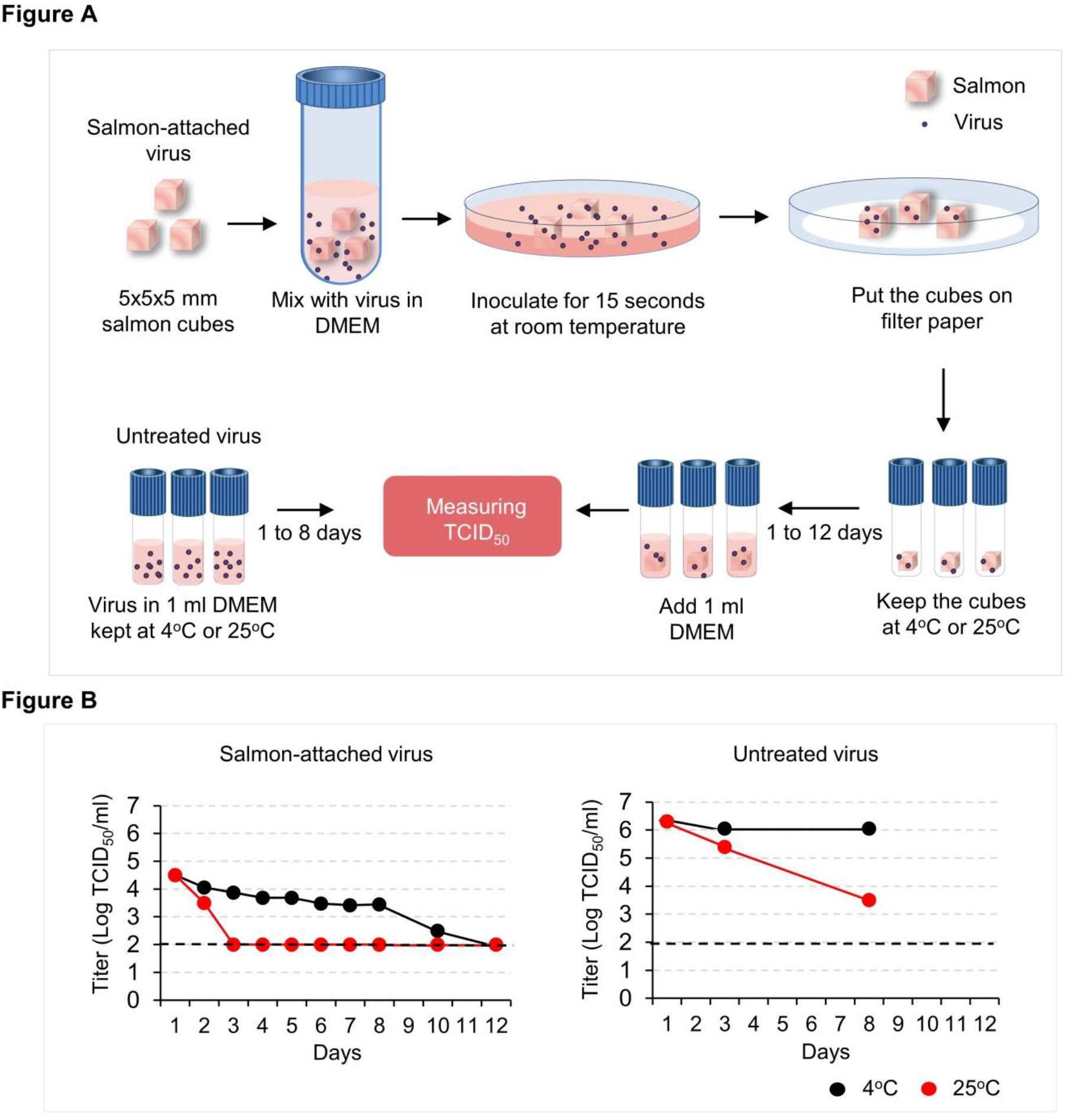
Viability of salmon-attached and untreated SARS-CoV-2 in culture medium at 4°C and 25°C. Panel A is the overview of the study design and experimental procedure. In panel B, the titer of SARS-CoV-2 was quantified by end-point titration on Vero E6 cells and is expressed as log_10_ TCID_50_ /mL. Plots show the means of data from two or three samples. The dashed lines indicate the limit of detection, which were 10^2^ TCID_50_ /mL.

Imported and exported fish must be transported under a low-temperature (e.g., 0 ~ 4°C) environment. Under such condition, SARS-CoV-2-contaminated fish from one country can be easily transported to another country within one week, thus serving as one of the sources for international transmission of SARS-CoV-2.

Different from vegetables and other food, fish have to be transported, stored and sold under a low-temperature environment. Fish are generally sold in quarters having temperatures much lower than regular room temperature. This means that virus attached on fish skin and sold in fish or seafood markets can survive for a long time.

In conclusion, fish-attached SARS-CoV-2 can survive for more than one week at 4°C, the temperature of refrigerators, cold rooms, or transport carriers for storage of fish before selling in the fish or seafood market. This calls for strict inspection or detection of SARS-CoV-2 as a critical new protocol in fish importation and exportation before allowing sales.

## Acknowledgments

This work was supported by the Key Research and Development Project of Guangdong Province (202020012624900001) and National Natural Science Foundation Grants (31830097 and 31802174). We thank Zhonghua Liu from Jiangsu Bioperfectus Technologies Co., Ltd. for his unconditional help while carrying out the experimental design for this work.

## About the Author

Dr. Dai is Associate Professor, South China Agricultural University. Her research interest is host antiviral immune response.

## Appendix

### Methods

#### Virus

SARS-CoV-2 GDPCC-nCOV4 strain used in this experiment was isolated and provided by Guangdong Provincial Center for Disease Control and Prevention. All work with SARS-CoV-2 was performed under BSL3 containment at the South China Agricultural University ABSL3 laboratory.

#### Titration of salmon-attached virus and untreated SARS-CoV-2

Salmon purchased from a salmon shop in Guangzhou was detected negative for SARS-CoV-2 with a 2019-nCoV detection kit (Bioperfectus, Taizhou, China) using real-time RT-PCR. Individual salmon cubes (5×5×5 mm) were placed in a 50 mL tube containing 13 mL liquid of SARS-CoV-2 at 3.16×10^6^ TCID_50_/mL, and the tube was gently inverted 5 times. The salmon cubes were transferred into the 10 cm dish and incubated for 15 seconds at room temperature and then put on filter paper in another 10 cm dish to remove the excess virus liquid. Salmon cubes were transferred to 1.5 mL freezing tubes and stored at 4°C and 25°C, respectively. On day 1, 2, 3, 4, 5, 6, 7, 8, 10, and 12, respectively, one freezing tube was taken out, to which 1 mL DMEM culture medium was added, oscillated for 5 seconds, and then centrifuged at 6,000 rpm for 5 minutes at 4°C. About 0.5 mL of the liquid was transferred to a new freezing tube and kept at −80°C until viral titration (Figure A). The untreated virus in culture medium was stored at 4°C and 25°C, respectively. On day 1, 3, and 8, 1 mL of the viral liquid was taken out and put in a freezing tube, which was kept at −80°C until viral titration (Figure A). The virus titer (50% tissue culture infectious dose, TCID_50_ per mL) was quantified by end-point titration assay on Vero E6 cells (*1*). The detection limit of the typical TCID_50_ assay used in this study was 10^2^ TCID_50_ / mL (*1*).

**Supplementary Table 1.**
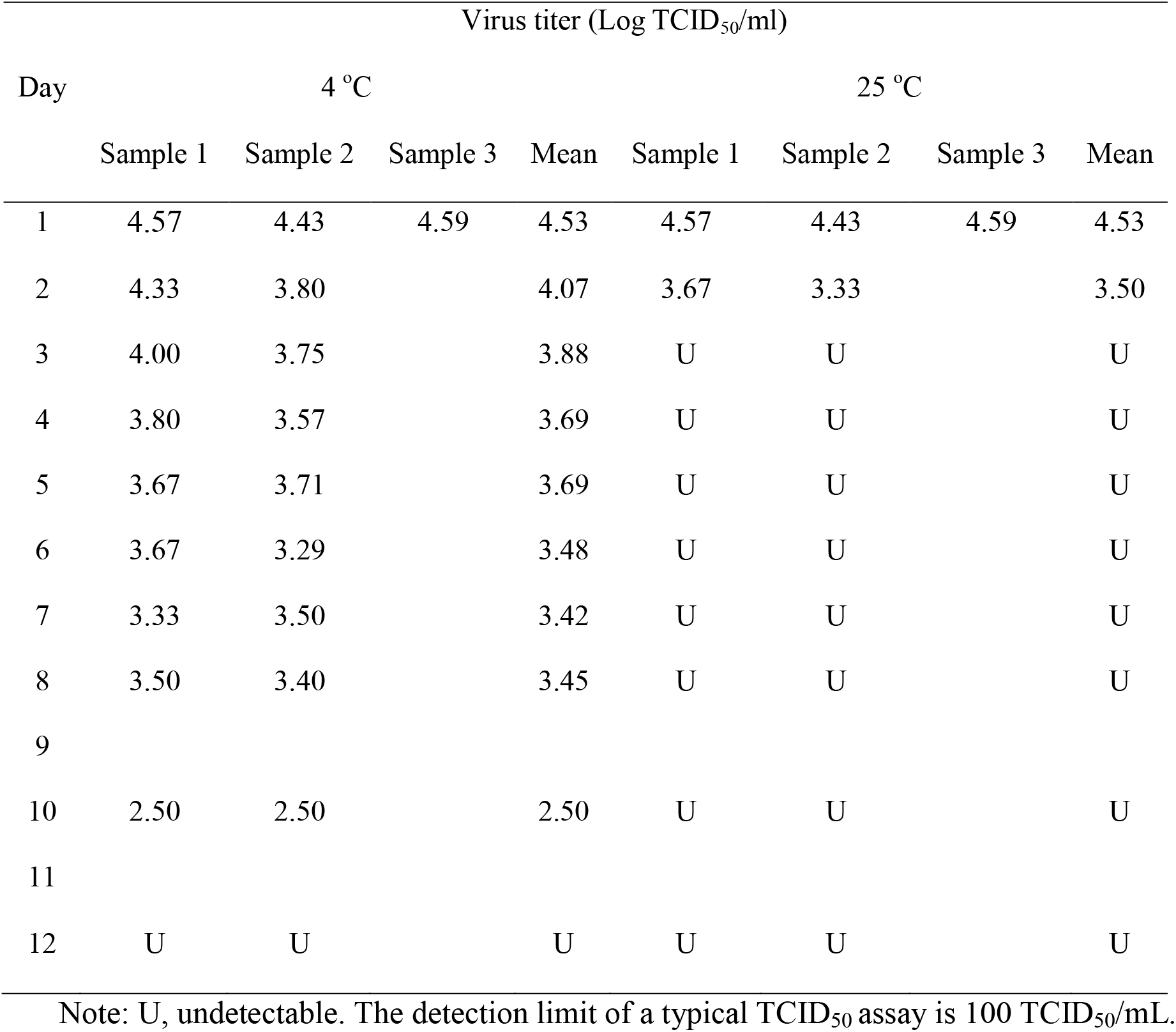
Stability of salmon-attached SARS-CoV-2 at 4°C and 25°C

**Supplementary Table 2.**
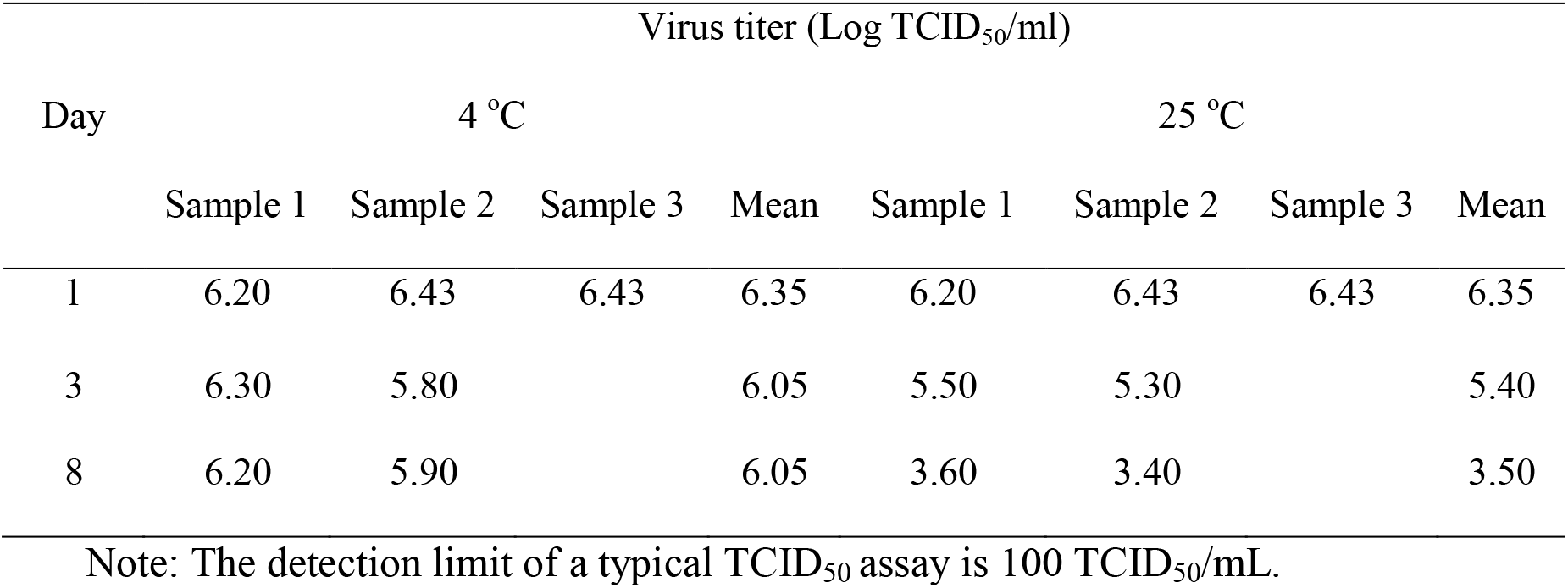
Stability of untreated SARS-CoV-2 in culture medium at 4°C and 25°C

## References

1. Normile D. Source of Beijing’s big new COVID-19 outbreak is still a mystery. https://www.sciencemag.org/news/2020/06/source-beijing-s-big-new-covid-19-outbreak-still-mystery (accessed September 5, 2020).

2. COVID-19 outbreaks in Wuhan, Beijing and Dalian share certain similarities: China’s top epidemiologist. https://www.globaltimes.cn/content/1196130.shtml (accessed September 5, 2020).

3. Waltenburg MA, Victoroff T, Rose CE, et al. Up.date: COVID-19 Among Workers in Meat and Poultry Processing Facilities - United States, April-May 2020. MMWR Morb Mortal Wkly Rep. 2020;69:887–92.

4. Dao D. COVID-19 infections force Thai Union to close fish canning factory in Ghana. https://www.seafoodsource.com/news/processing-equipment/covid-19-infections-force-thai-union-to-close-fish-factory-in-ghana (accessed September 5, 2020).

5. Coronavirus shutdown at Cedar Meats over as abattoir resumes full operations. https://www.abc.net.au/news/2020-05-27/coronavirus-shutdown-melbourne-cedar-meatsworkers-return/12289970 September 5, 2020).

6. Pan X, Ojcius DM, Gao T, Li Z, Pan C, Pan, C. Lessons learned from the 2019-nCoV epidemic on prevention of future infectious diseases. Microbes Infect, 2020;22:86–91.

7. Seeding of outbreaks of COVID-19 by contaminated fresh and frozen food. bioRvix. https://doi.org/10.1101/2020.08.17.255166.

8. van Doremalen N, Bushmaker T, Morris DH, et al. Aerosol and Surface Stability of SARS-CoV-2 as Compared with SARS-CoV-1. New Engl J Med. 2020;382:1564–7.

## References

1. van Doremalen N, Bushmaker T, Morris DH, et al. Aerosol and Surface Stability of SARS-CoV-2 as Compared with SARS-CoV-1. New Engl J Med. 2020;382:1564–7.

